# Ranking Cancer Drivers via Betweenness-based Outlier Detection and Random Walks

**DOI:** 10.1101/2020.03.03.974295

**Authors:** Cesim Erten, Aissa Houdjedj, Hilal Kazan

**Affiliations:** Department of Computer Engineering, Antalya Bilim University, Antalya, Turkey; Electrical and Computer Engineering Graduate Program, Antalya Bilim University, Antalya, Turkey

**Keywords:** Driver gene prioritization, bipartite graph, betweenness centrality, network diffusion

## Abstract

**Background:** Recent cancer genomic studies have generated detailed molecular data on a large number of cancer patients. A key remaining problem in cancer genomics is the identification of driver genes. Results: We propose BetweenNet, a computational approach that integrates genomic data with a protein-protein interaction network to identify cancer driver genes. BetweenNet utilizes a measure based on betweenness centrality on patient specific networks to identify the so-called *outlier genes* that correspond to dysregulated genes for each patient. Setting up the relationship between the mutated genes and the outliers through a bipartite graph, it employs a random-walk process on the graph, which provides the final prioritization of the mutated genes. We compare BetweenNet against state-of-the art cancer gene prioritization methods on lung, breast, and pan-cancer datasets. Conclusions: Our evaluations show that BetweenNet is better at recovering known cancer genes based on multiple reference databases. Additionally, we show that the GO terms and the reference pathways enriched in BetweenNet ranked genes and those that are enriched in known cancer genes overlap significantly when compared to the overlaps achieved by the rankings of the alternative methods.

## 1 Background

Cancer is a complex disease arising in many cases from the effects of multiple genetic changes that give rise to pathway dysregulation through alterations in copy number, DNA methylation, gene expression, and molecular function [1, 2]. Recent cancer genomics projects such as The Cancer Genome Atlas (TCGA) have created a comprehensive catalog of somatic mutations across all major cancer types. A key current challenge in cancer genomics is to distinguish driver mutations that are causal for cancer progression from passenger mutations that do not confer any selective advantage. Consequently, several computational methods have been proposed for the identification of cancer driver genes or driver modules of genes by integrating mutations data with various other types of genetic data [3, 4, 5, 6, 7, 8, 9, 10]; see [11, 12, 13, 14] for recent comprehensive evaluations and surveys on the topic. Rather than outputting a set of candidate driver genes or modules, a subclass of cancer driver identification methods output a prioritized list of genes ranked by their cancer driving potential. Early approaches in this group have utilized the mutation frequency of each gene by comparing with background mutation rates [15, 16, 17]. However with a careful review of the existing cancer catalogues it is easy to observe that most tumors share only a small portion of the set of all mutated genes, giving rise to the so called *tumor heterogeneity problem*; methods solely based on mutation rates suffer from low sensitivity due to the existence of long-tail of infrequently mutated genes [4, 18].

One strategy that aims to tackle the long-tail phenomenon is to move from a mutation-centric point of view to a *guilt by association* viewpoint where a correlation between differentially expressed genes and mutated genes are sought. This strategy assumes that even though different sets of genes are mutated in different patients, each of the candidate driver mutations tends to affect a large number of differentially expressed genes. Masica and Karchin present one of the early models based on such a strategy by employing statistical methods for setting up the correlation between mutated genes and the differentialy expressed genes to identify candidate drivers [1]. Many different models follow a similar trail by further incorporating biological pathway/network information for setting up such a correlation [6, 19, 20, 21, 22, 23]. DriverNet is among the notable approaches employing mutations data in addition to gene expression and biological network data [19]. It prioritizes mutated genes based on their degrees of network connectivity to dysregulated genes in tumor samples where dysregulation is determined via differential gene expression. Many subsequent approaches are inspired by DriverNet [20, 21, 22, 23]. Among them DawnRank [20], the algorithm by Shi et al. [21], and Subdyquency [23] employ, on top of the overall DriverNet model, versions of heat diffusion on the networks integrating data in the form of biological interactions, mutations, and gene expression. Heat diffusion is a technique employed commonly in many cancer driver gene or gene module discovery algorithms [9, 24, 25, 26, 27, 28]. It generally serves two purposes simultaneously. On the one hand, since the employed interactions data is usually erroneous, diffusing any type of information through the network of interactions, fixes any potential issues arising from missing links in the network. On the other hand, via the diffusion process, it is possible to observe the extent of an effect such as mutation frequency of a gene, at various distant loci in the network. LNDriver extends the DriverNet concept by taking into account gene lengths of the mutated genes to filter out genes that are mutated with high probability due to their lengths [22]. It should be noted that DawnRank and Subdyquency differ slightly from other approaches; the former can identify patient-specific candidate drivers and the latter employs subcellular localization information in addition to the data made of use in the other methods. There are other driver gene prioritization methods that deviate from the overall guilt by association framework, but nevertheless employ different types of genetic data together with the mutations data. IntDriver utilizes an interaction network and gene ontology data within a matrix factorization framework [29]. Dopazo and Erten employ paired data to generate tumor and normal interaction networks filtered with mutations and gene expression data, and measure the efficacy of various graph-theoretical measures in prioritizing breast cancer genes [6]. Note that among the discussed methods DawnRank also utilizes paired data, both from the tumor and the normal samples.

We propose BetweenNet algorithm for cancer driver gene prioritization. Similar to the methods proposed in [20, 21, 22, 23], BetweenNet is also inspired by the DriverNet framework in that it relates the mutated genes and the so-called *outlier genes* corresponding to the dysregulated genes in each patient through a bipartite influence graph. However different from DriverNet and the previous other methods based on it, BetweenNet determines outlier genes based on the betweenness centrality values of the genes in personalized networks. A second contribution of BetweenNet is the employment of a random-walk process on the resulting influence bipartite graph. Random-walks have been utilized in this context previously [21, 23]. However our application of random walk with restart on the whole influence graph is quite different from the two-step or three-step employment of the diffusion process on a per patient basis described in these methods. Through extensive evaluations we demonstrate that BetweenNet outperforms the alternative methods in recovering known reference genes and in providing functionally coherent rankings when compared to the enriched GO terms or the enriched known functional pathways.

## 2 Methods

We describe the details of the main steps of the BetweenNet algorithm in this section. Figure 1 provides an overview of the algorithm.

**Figure 1.**
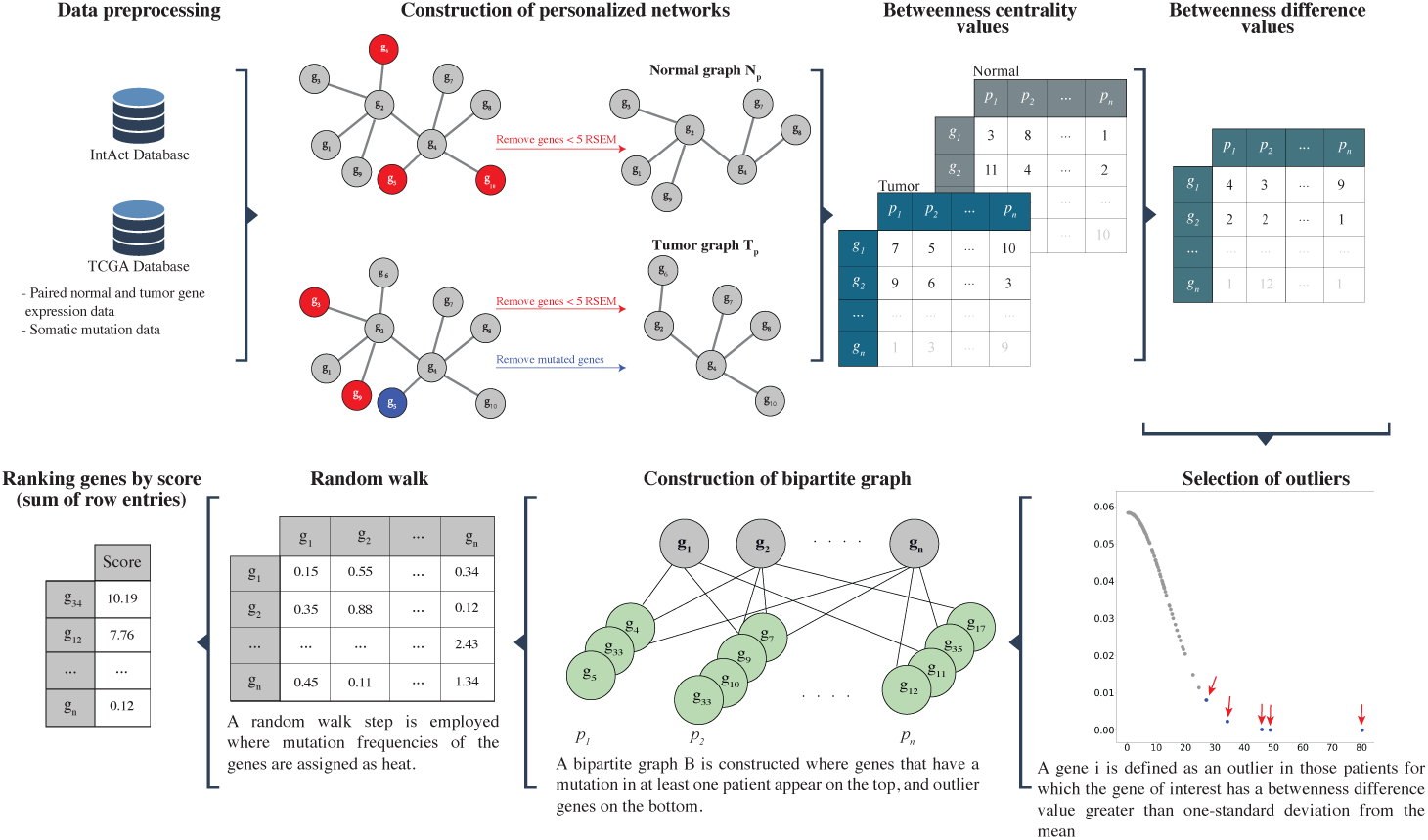
Main steps of the BetweenNet algorithm.

### 2.1 Input data sets and data preparation

In order to construct the pan-cancer cohort, we first identify the cancer types that have more than 10 paired measurements from normal and tumor samples in the TCGA cohort [30] (Supplementary Table 1). We then take the union of all the samples from these cancer types to form the cohort. In addition to the pan-cancer data, we perform separate evaluations on two cancer types. These are breast cancer (BRCA) with 112 samples, lung cancer (LUSC + LUAD) with 109 samples. We download the gene expression (RSEM normalized values [31]) and somatic mutation data for these patients from the Firebrowse database (*http://firebrowse.org*; version 2016 01 28). We exclude the silent mutations in the calculation of mutation frequencies. In addition to the gene expression and mutations data, we also employ protein-protein interactions data which we gather from the *H. Sapiens* PPI network of the IntAct database [32]. We preprocess the PPI network so that multiple ids for the same protein are merged to a single id. The resulting network contains 16,513 nodes and 189,520 edges.

### 2.2 Construction of personalized networks

Let *G* = (*V, E*) represent the reference *H. Sapiens* PPI network where each vertex *u*_*i*_ ∈ *V* denotes a gene *i* whose expression gives rise to the corresponding protein in the network. Each undirected edge (*u*_*i*_, *u*_*j*_) ∈ *E* denotes the interaction among the proteins corresponding to the genes *i, j*. Let *P* represent the set of patient samples. For each patient *p* ∈ *P*, we define two graphs *N*_*p*_ and *T*_*p*_ that represent the PPI networks of the normal and tumor samples, respectively. To construct *N*_*p*_, we start with the reference PPI network *G* and remove the nodes that correspond to the genes that are not expressed in the normal sample of the patient *p*. We deem genes with < 5 RSEM as not expressed. To construct *T*_*p*_, we remove two sets of genes: (i) genes with < 5 RSEM in the tumor sample; (ii) genes that contain non-silent mutations in the tumor sample.

### 2.3 Calculation of betweenness centrality values

The standard definition of the betweenness centrality ignores the length of a shortest path. Since considering very long paths as functional relations may not be biologically meaningful, we use a variant of the betweenness centrality called *k-betweenness*, where only shortest paths of length ≤ *k* are included in the calculations [33]. Given an unweighted graph *G* = (*V, E*), k-betweenness value of a node that corresponds to gene *i* is defined as follows:

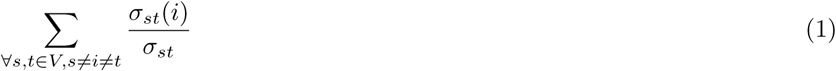

where *σ*_*st*_ is the number of shortest paths of length ≤ *k* between genes *s* and *t*, and *σ*_*st*_(*i*) is the number of such paths that pass through gene *i*. We choose *k* to be 3 in this study and utilize the algorithm presented in Brandes *et al.* to efficiently calculate the k-betweenness values [34]. Let 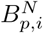 and 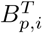 denote the 3-betweenness centrality values of the gene *i* in the *N*_*p*_ and *T*_*p*_ graphs of the patient *p*, respectively. We define 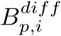 as 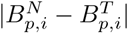.

### 2.4 Selection of outlier genes

For each gene *i*, we plot the 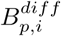 values across all the patients. We observe that the distribution can be approximated with a truncated normal distribution (Supplementary Figure 1). We use the *truncnorm* function in Python to estimate the mean and standard deviation of the distribution. A gene *i* is defined as an *outlier* in patient *p*, if 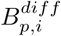 is greater than one standard deviation from the mean. We repeat this process for each gene and construct a set of outlier genes for each patient.

### 2.5 Construction of the bipartite graph

Similar to DriverNet, we construct a bipartite graph *B* that models the relationship between the set of mutated genes and the outliers. The *mutations partition* of the bipartite graph consists of the genes that have a mutation in at least one patient and the *outliers partition* consists of the outlier genes of all the patients in the cohort. Note that a gene *j* can be an outlier for multiple patients. In such a case, each occurence of a gene is represented with a distinct node in the outliers partition of *B*. Assuming *j* is an outlier gene for patient *p*, let 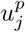 be the node corresponding to it in the outliers partition. For a node *u*_*i*_ in the mutations partition, edge (*u*_*i*_, 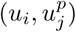) is inserted in *B*, if gene *i* is mutated in *p* and (*u*_*i*_, *u*_*j*_) is an edge in *G*.

### 2.6 Random walk on the bipartite graph

We apply a random walk on the bipartite graph *B*. The mutation frequencies of the genes are assigned as initial heat values to be diffused throughout the network during random walk. Let *MF* (*i*) denote the mutation frequency of gene *i*, that is, the number of patients where *i* has a non-silent mutation divided by the total number of patients. Note that heat values are assigned to genes on both sides of the bipartite graph. The random walk starts at a node *u*_*i*_ in *B* and at each time step moves to one of *u*_*i*_’s neighbors with probability 1 − *β* (0 ≤ *β* ≤ 1). The walk can also restart from *u*_*i*_ with probability *β*, called the *restart probability*. This process can be defined by a transition matrix *T* which is constructed by setting 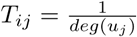 if (*u*_*i*_, *u*_*j*_) ∈ *E*, and *T*_*ij*_ = 0 otherwise. Here, *deg*(*u*_*j*_) corresponds to the degree of the node *u*_*j*_. Thus *T*_*ij*_ can be interpreted as the probability that a simple random walk will transition from *u*_*j*_ to *u*_*i*_. The random walk process can also be considered as a network propagation process by the equation, *F*_*t*+1_ = (1 − *β*)*TF*_*t*_ + *βF*_0_, where *F*_*t*_ is the distribution of walkers after *t* steps and *F*_0_ is the diagonal matrix with initial heat values, that is *F*_0_[*i, i*] = *MF* (*i*). We compute the final distribution of the walk by calculating the *F* matrix iteratively until convergence. We set *β* = 0.4 as in previous studies where the effect of varying *β* has been found to be not significant [9, 28]. After convergence, genes are prioritized by the sum of incoming edge weights.

### 2.7 Compiling reference gene sets

We compile known cancer genes from the databases Cancer Gene Census (CGC) [35] and CancerMine [36]. From the former, we obtain the list of 728 genes that are found to be associated with cancer. CancerMine uses text-mining to catalogue cancer associated genes where it also extracts information about the type of the cancer. We compile two lists of genes that have at least 3 and 5 citations, respectively. Hereafter, these two reference gene sets are named *CancerMine3* and *CancerMine5*. The number of genes in *CancerMine3* and *CancerMine5* reference sets for each cancer type (i.e., lung, breast) and for pan-cancer cohort are available in the Supplementary Tables 2-4. For lung cancer, we are unable to use *CancerMine5* as a reference due to its small size.

### 2.8 Enrichment analysis with Gene Ontology and pathway databases

For Gene Ontology (GO) [37] term analysis, we use *goatools*. We download go-basic.obo file from http://geneontology.org/docs/download-ontology/ on June 26th of 2019. We restrict the gene annotations to level 5 by ignoring the higher-level annotations and replacing the deeper-level category annotations with their ancestors at the restricted level.

For the pathway analysis, we use the *AllEnricher* tool with Reactome and Kyoto Encyclopedia of Genes and Genomes (KEGG) [38] pathways. Both *goatools* and *AllEnricher* use Fisher’s exact test to calculate p-values and False Discovery Rate (FDR) for multiple testing correction. We use 0.05 as the p-value cutoff to determine significant enrichments.

## 3 Results

We implemented the betweenness centrality measurement algorithm in C++ using the *LEDA* (Library of Efficient Data types and Algorithms) library. The remaining steps are implemented in Python using *NetworkX* library. All the code and necessary datasets are available at https://github.com/abu-compbio/BetweenNET. We compare BetweenNet results against those of five other existing cancer driver prioritization methods: DriverNet, Subdyquency, DawnRank, IntDriver, and Dopazo and Erten’s prioritization method based on betweenness centrality values, hereafter named only *Betweenness*. Note that for the Betweenness method, although the original method ranks all genes, here we only rank mutated genes using the same method for a fair comparison, since all the other methods under consideration are designed to rank mutated genes only. DriverNet is chosen due to its close connection to our work. DawnRank and Subdyquency are included as they extend and improve over DriverNet. Betweenness is included as a baseline since our method utilizes a variation of betweenness differences in identifying outlier genes. Finally, IntDriver is included to represent the performance of a distinct strategy that is based on matrix factorization. We evaluate the methods with three datasets: lung cancer, breast cancer, and pan-cancer samples.

### 3.1 Evaluations with respect to reference cancer gene sets

We first compare the methods based on their ability to recover the sets of known cancer genes. For this, we compute true positive and false positive rates for the top 1000 genes and calculate the area under the ROC (AUROC). Figure 2 shows the ROCs obtained from lung cancer data. In Figure 2-a all CGC genes are used as reference, whereas in Figure 2-b genes with mutation frequencies ≤ 2%, namely the *rare drivers*, are included. Figure 2-c is obtained with CancerMine3 as the reference set. BetweenNet achieves a higher AUROC value than all the alternatives for all three reference sets, though the improved performance is more prominent for the first two reference sets. With the CGC reference set, the second ranked method is Subdyquency. This is followed by Betweenness and DawnRank with similar performance with respect to each other. On the other hand, for the CGC-rare reference set, Betweenness and DawnRank perform considerably better than the other three methods. DawnRank’s good performance is in line with its focus on finding rare drivers by identifying drivers in a personalized fashion rather than using the mutation frequency of a gene across all the patients. Interestingly, Betweenness results in poor performance with the CancerMine3 reference set where it ranks the second worst method. For all three reference sets, the IntDriver method performs the worst. Overall, these results illustrate the superiority of BetweenNet as it can find both rare and common drivers more accurately.

**Figure 2.**
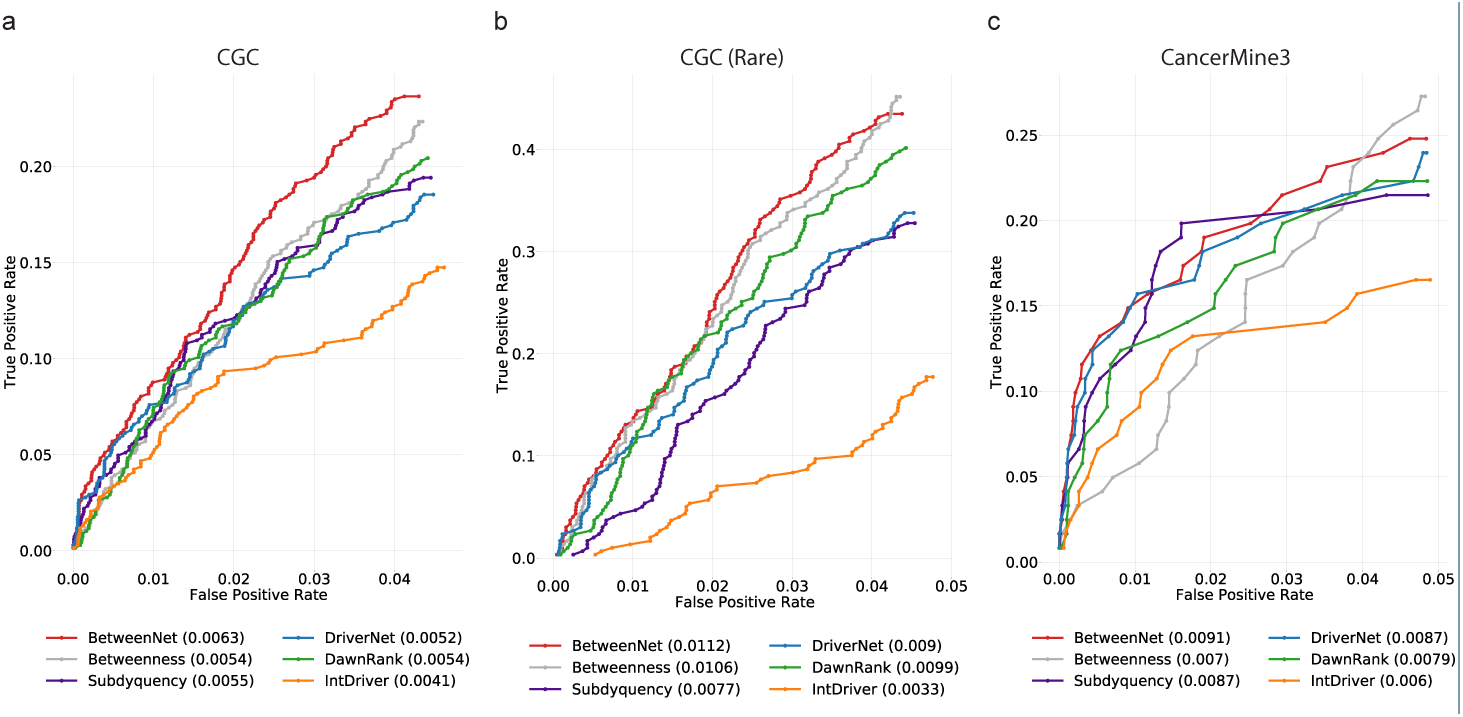
The fraction of recovered reference genes is shown with a ROC curve for lung cancer data a) *CGC* genes are used as reference. b) *CGC* rare genes are used as reference. c) *CancerMine3* genes are used as reference.

Figure 3 depicts analogous results for the breast cancer data. As before, BetweenNet achieves the top performance with all three reference sets. For the CGC reference set, Subdyquency and DriverNet rank the second and third, respectively. Betweenness and DawnRank have a similar performance which is worse than Subdyquency and DriverNet. Finally, IntDriver provides the worst performance. For CGC-rare we see a similar performance of the methods except that Betweenness ranks third after BetweenNet and Subdquency. For CancerMine3, BetweenNet ranks the best which is followed by DawnRank and Betweenness. Results with respect to the CancerMine5 reference set are quite similar and are available in the Supplementary Figure 2-a.

**Figure 3.**
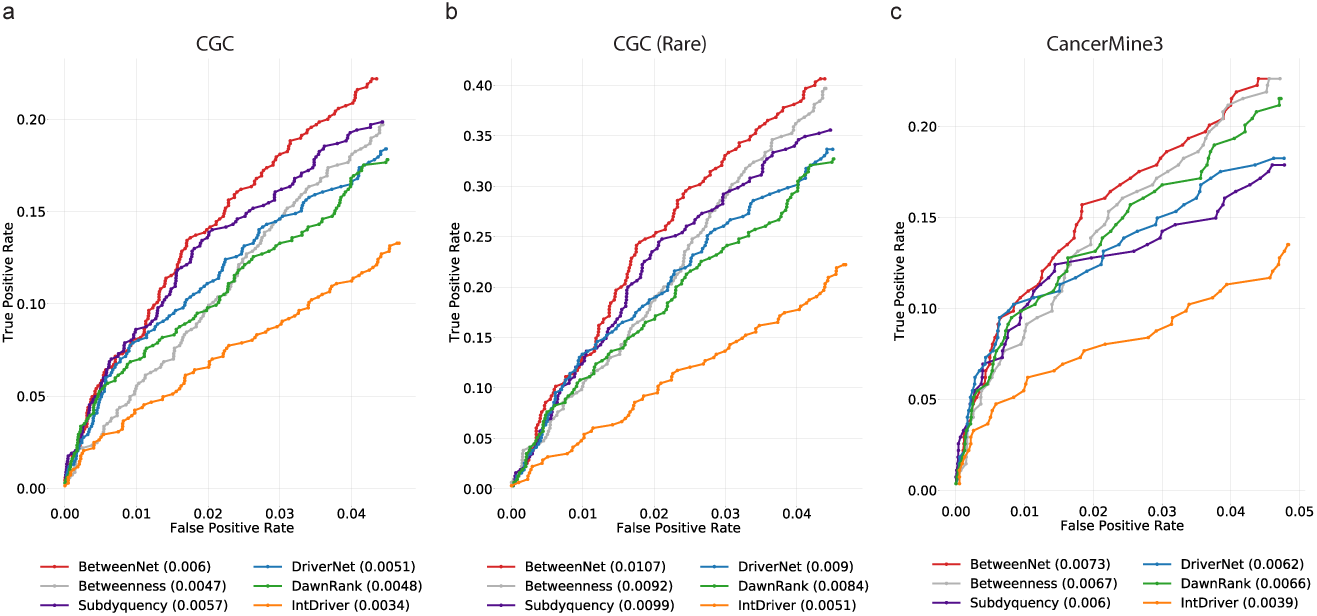
The fraction of recovered reference genes is shown with a ROC curve for breast cancer data a) *CGC* genes are used as reference. b) *CGC* rare genes are used as reference. c) *CancerMine3* genes are used as reference.

Lastly, Figure 4 shows the results with respect to the pan-cancer dataset. BetweenNet’s gene ranking results in the highest AUROC in all three cases, though the difference between BetweenNet and the second best performing method is more prominent for the CancerMine3 reference set. For the CGC reference gene set, BetweenNet, Subdquency, and DriverNet perform significantly better than the other three methods. For the CGC-rare reference set, DawnRank’s performance gets much closer to that of the leading methods, as we previously observe for the other datasets. For the CancerMine3 set, DriverNet and Betweenness can be considered as the second ranking group of methods, whereas DawnRank and IntDriver perform significantly worse than the others. Results with CancerMine5 are similar to those of CancerMine3 and are available in the Supplementary Figure 2-b.

**Figure 4.**
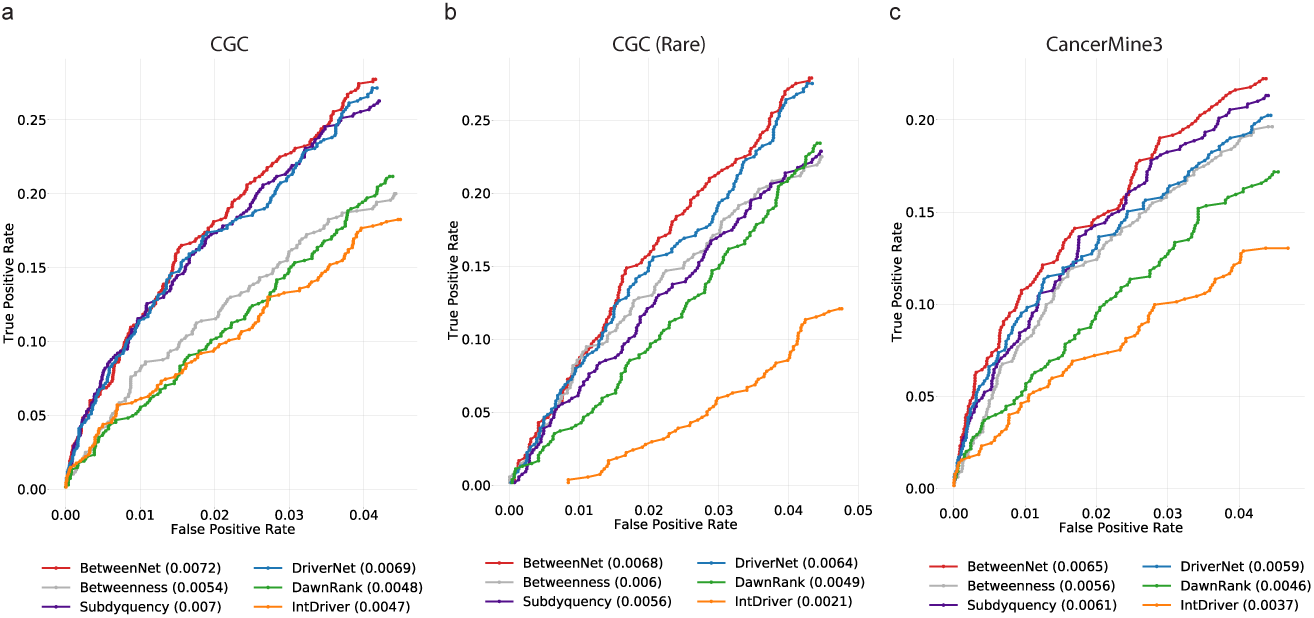
The fraction of recovered reference genes is shown with a ROC plot for pan-cancer data a) *CGC* genes are used as reference. b) *CGC* rare genes are used as reference. c) *CancerMine3* genes are used as reference.

### 3.2 Evaluations based on functional and pathway analysis

Reference cancer driver gene sets might be incomplete and biased. As such, rather than only finding exact matches between the output gene sets and the reference gene sets, we also define other metrics that measure how well the associated functions of the genes of the two sets match. One such metric is based on GO consistency (GOC) and the other is based on pathway information. For the former, we find the GO terms enriched in the output gene sets and in the reference gene sets, and check whether the corresponding GO terms overlap. The underlying assumption is that the reference cancer genes and the predicted cancer genes should have similar biological functions. We find the enriched GO terms in the ranked gene sets of varying total sizes from 100 to 500 in the increments of 100 for each method under consideration. We repeat the same GO term enrichment analysis with the reference gene set. We then compute the GOC value between the enriched GO terms of the ranked gene set and those of the reference set, which is defined as the ratio between the size of the intersection of the two sets and the size of the union [39]. Figure 5 shows the GOC values calculated for each cancer type and pan-cancer cohort. We observe that BetweenNet ranked genes perform the best for almost all total size values. For lung cancer, BetweenNet ranks the best in three out of five total size values, whereas Betweenness performs the best in the remaining two cases. DriverNet and Subdquency rank the third and fourth, respectively. This is followed by DawnRank and IntDriver with notably large differences between the performances of these two methods. For breast cancer, BetweenNet performs significantly better than the other methods. All the rest of the methods except IntDriver exhibit similar performances whereas IntDriver performs significantly worse than the others. For pan-cancer data, the methods can be ranked as follows: BetweenNet, DriverNet, Betweenness, Subdyquency, DawnRank, and IntDriver.

**Figure 5.**
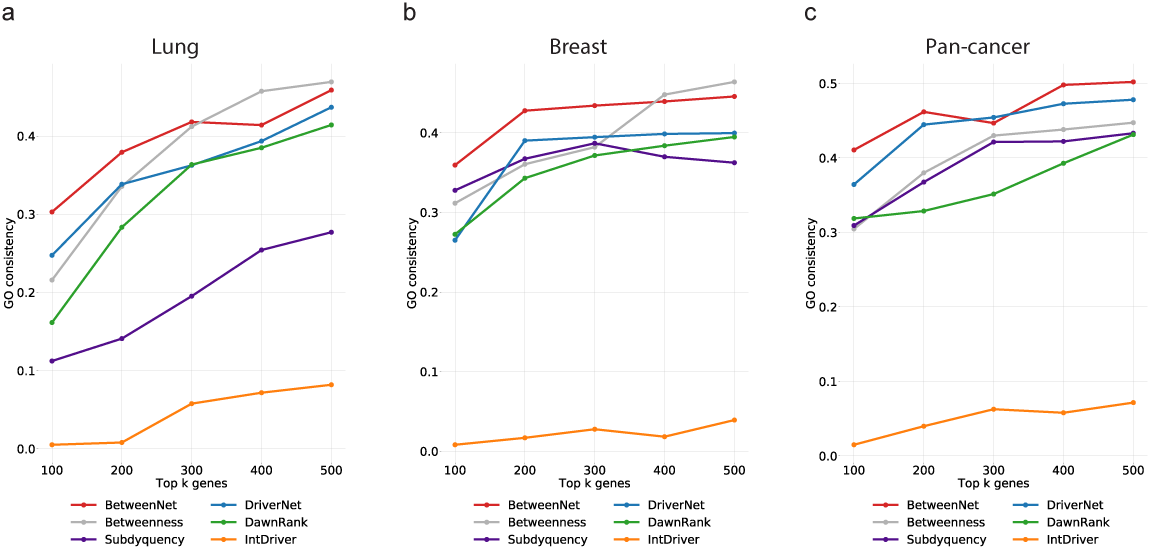
GO consistency values for a) lung cancer b) breast cancer c) pan-cancer cohort.

We repeat the same type of analysis with pathways as well, this time replacing GO term enrichment with pathway enrichment. Namely, we identify the pathways enriched in the reference set of genes and the set of genes output by a ranking method. We then compute the number of pathways common in both of these sets. Figure 6 shows the results with Reactome reference pathways for all cancer types. For all cases, BetweenNet outperforms the other methods with a large margin. For lung cancer, the second best method is Betweenness, whereas for breast and pan-cancer it is difficult to identify the second best method as it varies for each total size value. For breast and pan-cancer, IntDriver performs worse than the other methods. Results obtained with KEGG look similar and can be found in the supplementary.

**Figure 6.**
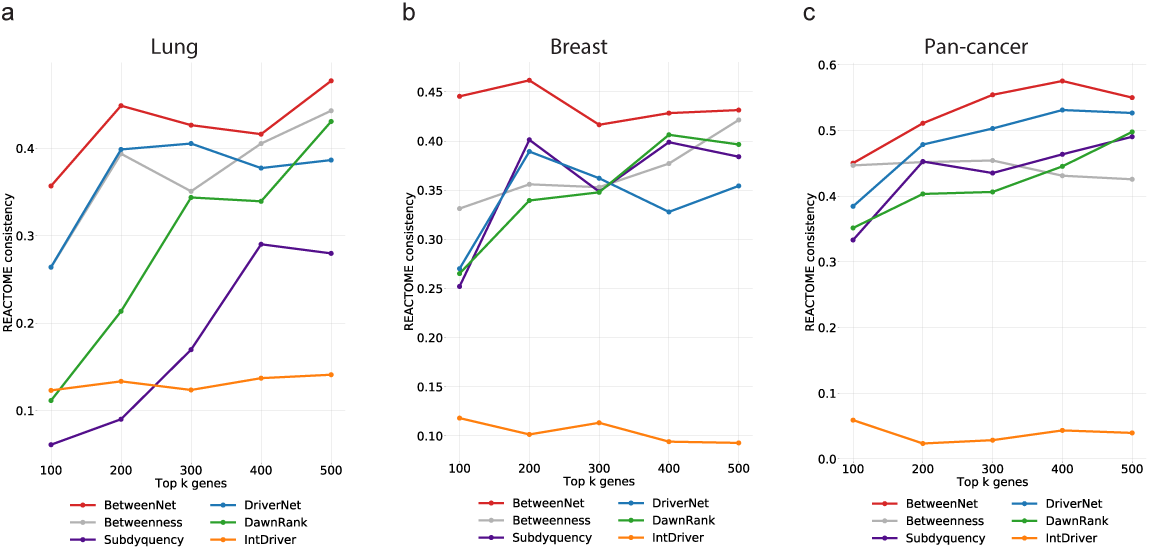
Reactome pathway consistency values for a) lung cancer b) breast cancer c) pan-cancer cohort.

## Analysis of BetweenNet ranked genes

We further explore BetweenNet’s top 20 ranking genes for each cancer type and identify those that do not appear in the cancer reference gene set CGC. We investigate the literature for existing work that study these genes within the context of cancer biology. For lung cancer, the gene ranked the sixth by BetweenNet, the Vascular cell adhesion molecule-1 (VCAM-1) gene has been found to have higher expression in cancer-associated fibroblasts and blocking it attenuates the proliferation and invasion of lung cancer cells [40]. VCAM-1 can easily be missed by the models solely based on mutation frequency, as it is mutated in only three patients across the lung cancer cohort. Indeed, only BetweenNet ranks this gene within the top 20. Another gene that ranks in our top 20 for lung cancer is Ryanodine receptor 2 (RYR2). This gene is found to be strongly associated with a subtype of lung adenocarcinoma that shows high tumor mutational burden [41]. RYR2 mutations were also found to be significantly higher in non-small cell lung cancer patients that live in highly polluted regions as compared to control regions [42]. Lastly, TP53BP1 is another gene that is only identified by BetweenNet within the top 20. TP53BP1 is a critical component of DNA damage response (DDR) machinery and defects of this machinery are involved in multiple processes of tumorigenesis [43, 44, 45]. A recent study showed that TP53BP1 protein expression is significantly reduced in several cancer tissues including lung carcinoma suggesting a role as tumor suppressor [46]. FN1 is within the top 20 genes of the Betwenness ranking for breast cancer. A recent study confirms that increased fibronectin is linked to metastasis through its role in facilitation of cancer cell attachment and spreading in 3D suspension culture of MDA-MB-231 cell line [47]. As such, existing studies indicate that FN1 is not a driver but play critical roles in metastasis which can still be of interest for therapeutic purposes. Another top ranking gene for breast cancer is GABARAP. GABARAP has been shown to involve in tumorigenesis in vivo by delaying cell death and its associated immune-related response [48].

In pan-cancer data, CDC5L, VCAM1, TP53BP1, FN1, MKI67, and PLEC are among the top 20 ranking genes that do not appear in CGC. For each of these genes, there are multiple studies in the literature indicating roles for cancer development and progression. Additionally, many of these genes also appear in top 20 for individual cancer types we study, that is in lung and breast cancer. For instance, CDC5L is a transcription factor that is associated with tumorigenesis in multiple cancer types including prostate, hepatocellular, colorectal, and bladder cancer [49, 50, 51, 52]. VCAM1 is linked to tumor immune evasion in more than ten cancer types [53, 54, 55, 56, 57, 58, 59, 60], TP53BP1 also appears in our *CancerMine3* list and its role in cancer development is recently reviewed in [61]. There are studies showing FN1’s aberrant expression in multiple cancer types [62, 63, 64, 65]. Lastly, MKI67 expression has been shown as a biomarker for multiple cancer types [66, 67, 68, 69, 70].

## Discussion

Having shown that BetweenNet performs better than existing methods, we finally investigate whether all components of BetweenNet contribute to this performance by testing against simpler alternatives. We test a version of BetweenNet where the random walk step is omitted. In this version, there are no edge weights and ranking is simply based on the degrees of the bipartite influence graph nodes corresponding to the mutated genes. We also include DriverNet in this comparison to evaluate the effects of the outlier selection process. Comparing the version of BetweenNet without the random walk step against DriverNet reveals the added value of defining outlier genes based on betweenness centrality values rather than gene expression. Supplementary Figures 4-6 compare these three models on lung, breast, and pan-cancer data. Except for experiments with *CancerMine3* in lung cancer and *CGC* in pan-cancer, we observe that BetweenNet achieves the top performance for all cancer types and for all reference gene sets.

## Conclusions

We propose BetweenNet, a novel cancer driver gene prioritization approach that integrates genomic data with the connectivity within PPI networks. One contribution of BetweenNet is the identification of patient specific dysregulated genes with a measure based on betweenness centrality on personalized networks. BetweenNet ranks mutated genes by their effects on dysregulated genes. To characterize these effects, a bipartite influence graph is formed to represent the relations between the mutated genes and dysregulated genes in each patient. Another contribution of BetweenNet is the employment of a random-walk process on the resulting influence bipartite graph. Through careful comparisons, we show that both the use of betweenness centrality metric and the employment of random walk have added values in identification of cancer driver genes. We also demonstrate that BetweenNet out-performs the alternative methods in recovering known reference genes and in providing functionally coherent rankings with three large-scale TCGA datasets: lung cancer, breast cancer, and pan-cancer samples. Additionally, we find that many of our top ranking genes that do not appear in reference cancer gene sets have roles in cancer development based on existing literature. Taken together, our results indicate that BetweenNet effectively integrates genomic data and connectivity information to prioritize cancer driver genes.

## Abbreviations

GO: Gene Ontology,
CGC: Cancer Gene Census,
TCGA: The Cancer Genome Atlas,
PPI: Protein-Protein Interaction,
BRCA: Breast Invasive Carcinoma
LUAD: Lung Adenocarcinoma,
LUSC: Lung Squamous Cell Carcinoma,
LEDA: Library of Efficient Data Types and Algorithms,
AUROC: Area Under the Receiver Operating Characteristics,
GOC: GO consistency.

## Ethics approval and consent to participate

Not applicable.

## Consent for publication

Not applicable.

## Availability of data and materials

The source code and datasets used in this research can be downloaded from https://github.com/abu-compbio/BetweenNET

## Authors’ contributions

Authors are listed in alphabetical order with respect to last names. CE and HK designed the overall framework. CE, AH, and HK contributed equally in the algorithm design. CE and HK designed the evaluation framework. AH implemented the algorithm, the evaluation scripts, and the performance tests. CE, AH, and HK contributed equally in writing the manuscript.

## Acknowledgements

Not applicable.

