## Supplementary for "Ranking Cancer Drivers via Betweenness-based Outlier Detection and Random Walks"

**Table 1 Pan-cancer size**

| Cancer type | Samples |
|---|---|
| STAD | 32 |
| LUSC | 51 |
| KIRC | 72 |
| KICH | 25 |
| KIRP | 32 |
| LIHC | 50 |
| LUAD | 58 |
| BRCA | 112 |
| ESCA | 11 |
| PRAD | 52 |
| HNSC | 43 |
| KICH | 25 |
| THCA | 58 |
| COADREAD | 32 |
| STES | 43 |
| BLCA | 19 |

**Table 2 Lung**

| Datasets | Genes | # of enriched GO terms | # of enriched Reactome pathways | # of enriched KEGG pathways |
|---|---|---|---|---|
| CGC | 723 | 1353 | 564 | 144 |
| CancerMine3 | 121 | 296 | 224 | 124 |
| CancerMine5 | 49 | - | - | - |

**Table 3 Breast**

| Datasets | Genes | # of enriched GO terms | # of enriched Reactome pathways | # of enriched KEGG pathways |
|---|---|---|---|---|
| CGC | 723 | 1353 | 564 | 144 |
| CancerMine3 | 274 | 1340 | 463 | 123 |
| CancerMine5 | 129 | 1049 | 337 | 111 |

**Table 4 Pan-cancer**

| Datasets | Genes | # of enriched GO terms | # of enriched Reactome pathways | # of enriched KEGG pathways |
|---|---|---|---|---|
| CGC | 723 | 1353 | 564 | 144 |
| CancerMine3 | 652 | 1107 | 471 | 124 |
| CancerMine5 | 337 | 1075 | 441 | 124 |

### Figures

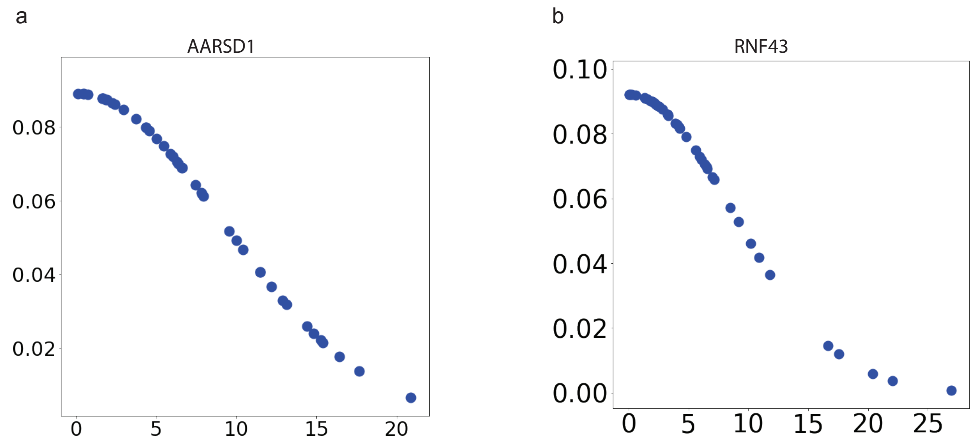


**Figure 1** The distribution of betweenness difference values for two selected genes AARSD1 and RNF43.

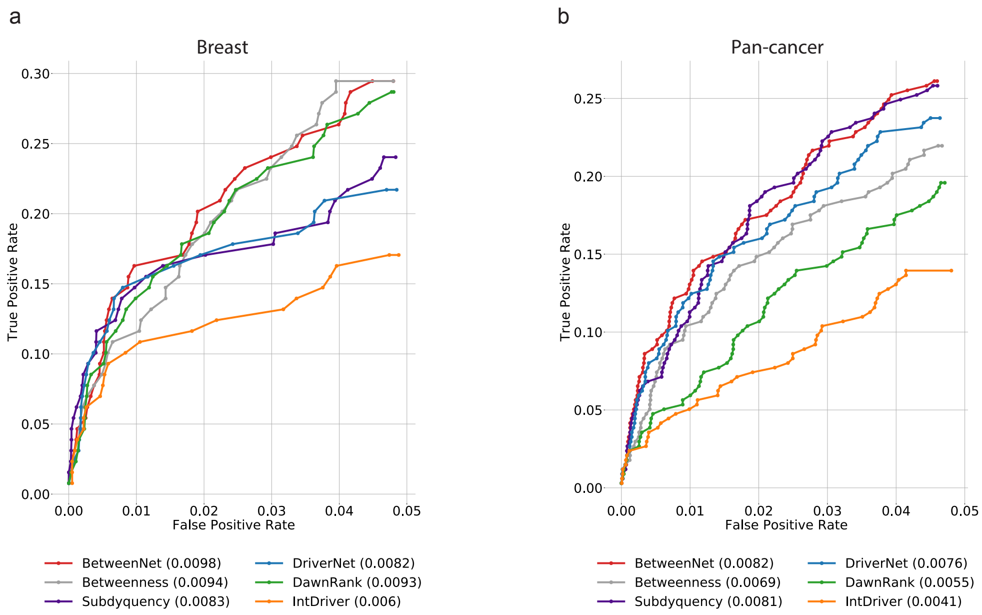


**Figure 2** The fraction of recovered reference genes is shown with a ROC curve for breast cancer data where *CancerMine5* genes are used as reference.

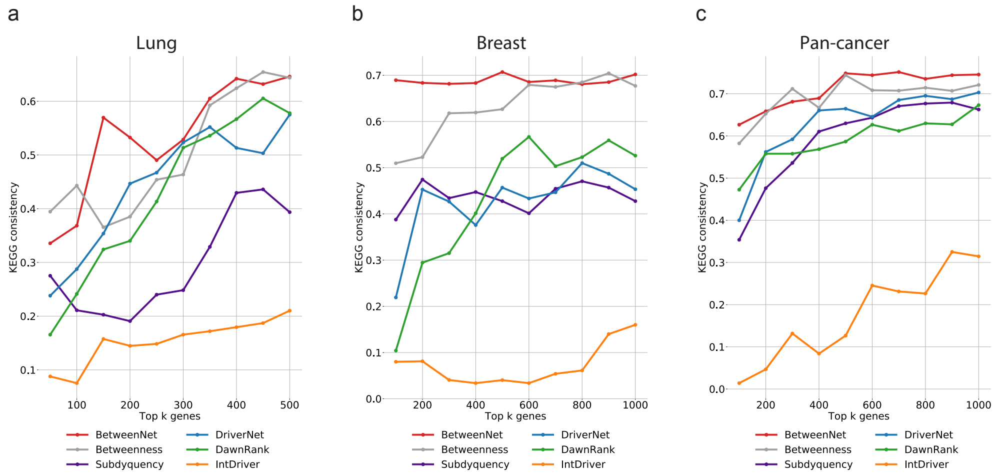


**Figure 3** KEGG pathway consistency values for a) lung cancer b) breast cancer c) pan-cancer cohort.

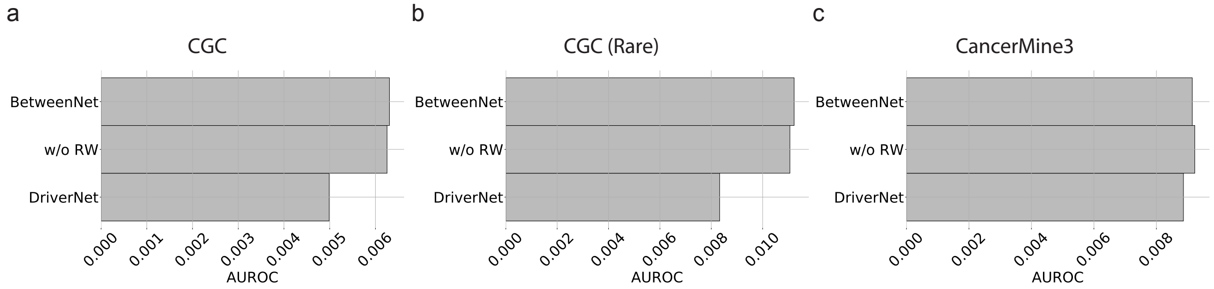


**Figure 4** Comparison of AUROC values of DriverNet, BetweenNet and BetweenNet's modified version where the random walk step is omitted for lung cancer. *CGC*, *CGC(Rare)*, *CancerMine3* genes are used as reference.

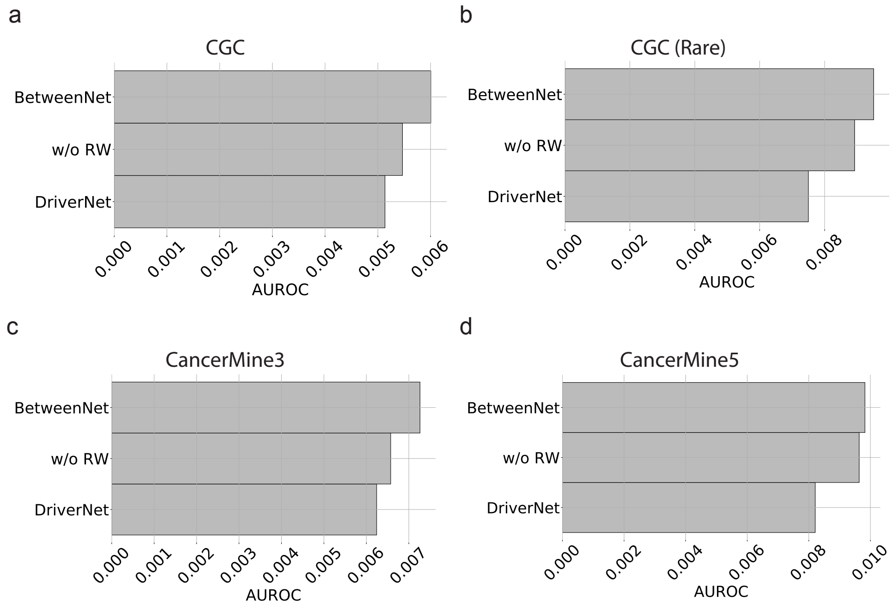


**Figure 5** Comparison of AUROC values of DriverNet, BetweenNet and BetweenNet's modified version where the random walk step is omitted for breast cancer data. *CGC*, *CGC(Rare)*, *CancerMine3* and *CancerMine5* genes are used as reference.

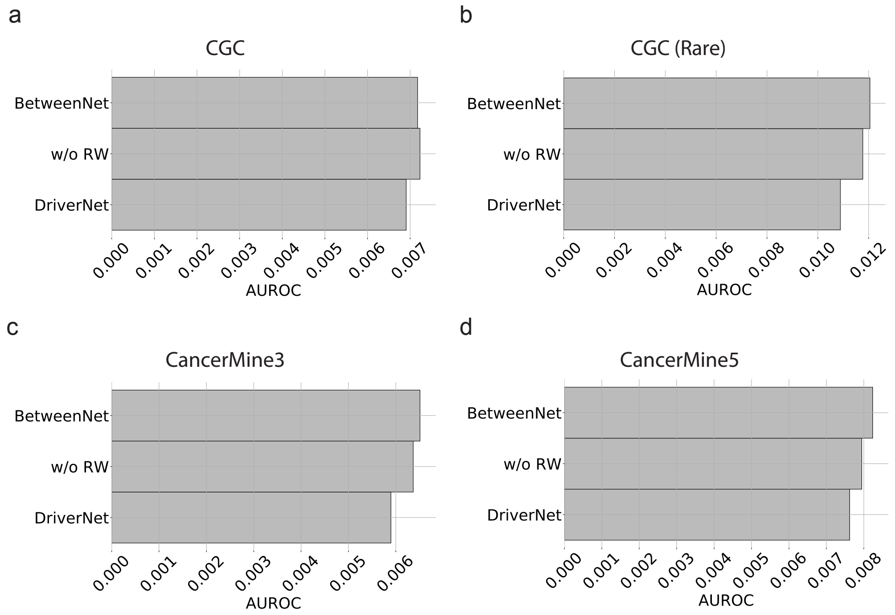


**Figure 6** *CGC*, *CGC(Rare)*, *CancerMine3* and *CancerMine5* genes are used as reference.
